# Combining AFM, XPS and chemical hydrolysis to understand the complexity and dynamics of *C. vulgaris* cell wall composition and architecture

**DOI:** 10.1101/2022.07.11.499560

**Authors:** Irem Demir-Yilmaz, Marion Schiavone, Jérôme Esvan, Pascal Guiraud, Cécile Formosa-Dague

## Abstract

The microalgae cell wall represents its interface with its environment and a strong barrier to disrupt in order to extract the cell’s products. Understanding its composition and architecture is a challenge that if overcome, could lead to substantial advancements in optimizing microalgae-production systems. However, the cell wall is a dynamic and complex structure that evolves depending on the growth phase or culture conditions. To apprehend this complexity, an experimental approach combining AFM, XPS, and chemical hydrolysis followed by HPAEC-PAD was developed to understand the cell wall of *Chlorella vulgaris*, a biotechnologically-relevant green microalgae species. Exponential and stationary growth stages were investigated, as well as saline stress condition inducing lipid production. Results showed that both the cell wall composition and architecture changes in stationary phase, with an increase of the lipidic fraction at the expanse of the proteic fraction, changes in the polysaccharidic composition, and a significant increase of its rigidity. Under saline stress, cell wall architecture seems to be affected as its rigidity decreases importantly. Altogether, this study demonstrates the power of combining these three techniques to give new insights into *C. vulgaris* cell wall, in terms of composition and architecture, and of its dynamics in different conditions.

---

Microalgae are unique microorganisms that convert light energy, water, and inorganic nutrients into a biomass resource rich in value-added products such as carbohydrates, proteins, or pigments [1]. In addition, microalgae are also highlighted as an alternative and renewable source of energy because of their important capacity to produce oil [1] that can be transformed into biofuel [2]. For this reason notably, a lot of attention has been paid to optimize culture conditions where the production of lipids by microalgae is maximized. For example, environmental stresses such as salinity increase, has been described to change the biomass composition and induce lipid accumulation in cells. For this reason, applying this type of stress, in different cultivation systems, has attracted a lot of interest for microalgae-based biofuel production [3]. In fact, high extracellular concentrations of Na^+^ directly influence the ionic balance inside cells and subsequently the cellular activities [4]. In particular, salinity stimulates the synthesis of storage neutral lipids, notably triacylglycerides (TAGs), produced as secondary metabolites and stored as energy reservoirs [5]. Then, in biofuel production systems, after cell harvesting, the lipids produced need to be extracted from cells, which is still a critical challenge that needs to be overcome by the industry and research community. For the moment, existing extraction techniques require a significant amount of chemicals or energy because of the chemically complex and structurally strong nature of microalgae cell walls [6]. Therefore, a better understanding of the ultrastructure and the composition of microalgae cell walls and their dynamics in different culture conditions, such as saline stress conditions, is needed to develop efficient and targeted extraction procedures of valuable intracellular products such as lipids.

The microalgae cell wall is a sophisticated structure, rigid and mechanically strong, which protects microalgae cells from predators and harsh environments [6]. In addition, the cell wall regulates the biological and biomechanical stability of the cell, influencing significantly its interaction with its surroundings [7]. But because of the diversity existing in cell wall composition and structure depending on microalgae strains, and because of its dynamics depending on culture conditions, the microalgae cell wall remains poorly understood. Yet, several studies have showed the development of methods to isolate microalgae cell walls and determine their composition. Most of these studies focus on determining the polysaccharidic composition of the cell wall, using a combination of mechanical disruption with chemical and/or enzymatic hydrolysis. For example, some studies applied mechanical disruption of cells to extract the cell walls, which were then treated using chemical treatments, with LiCl [8] or with strong acids [9], or/and using enzymatic hydrolysis [10]. This way the authors hydrolyze the polysaccharides present in the cell wall, and can then identify the saccharidic monomers released using high performance liquid chromatography (HPLC) [9,10]. Using these methods, Canelli *et al.* reported differences in the cell wall saccharidic composition of cells in exponential phase and in stationary phase for *C. vulgaris* [9], meaning that the amount and profile of sugars of the cell wall of microalgae depend on the growth stage, thus highlighting the dynamics of microalgae cell wall composition. To determine the total and relative composition of microalgae cell walls (proteic, polysaccharidic, and lipidic fractions), other studies use surface characterization techniques such as Fourier-transform Infrared spectroscopy (FTIR) [11] or cryo-X-ray Photoelectron Spectroscopy (cryo-XPS) [12,13]. For instance, a recent study by Shchukarev *et al.* could determine the surface composition in terms of polysaccharides, lipids and proteins of three different microalgae species, *C. vulgaris, Coelastrella sp.* and *S. obliquus* [13]. This study showed that the relative amounts of the different fractions were different for the three species considered, showing thus the diversity but most importantly the complexity of microalgae cell walls.

However, while these techniques, HPLC-based polysaccharidic determination and XPS, have been used separately so far to study microalgae cell walls, their combination could result in a more complete understanding of the composition and dynamic characterization of microalgae cell walls. In addition, other surface characterization techniques could be used to probe microalgae cell walls, such as atomic force microscopy (AFM). AFM, first developed in 1986 [14], is a powerful tool to image cells with nanometer-scale resolution and probe their nanomechanical properties under liquid conditions. For cell wall characterization, such technique has been used notably with yeast cells to measure both the roughness and the rigidity of the cell wall; such measurements brought valuable information to understand the architecture of cell walls [15]. In the microalgae field, AFM has also proven to be an efficient technique to understand microalgae cells, their morphology, their nanomechanical properties, and their response to different conditions such as environmental stress [16].

Among the wide variety of microalgae species, several have been considered for biofuel production, including *Chlorella vulgaris. C. vulgaris*, first discovered in 1890 by a Dutch researcher [17], is one of the most studied microalgae species mainly because of its biotechnological importance for the production of proteins used in nutrition and for biofuel production [18]. Indeed, *C. vulgaris* is a unicellular freshwater microalgae species able to accumulate significant amounts of lipids under specific culture conditions, with a fatty acid profile adapted for biofuel production [19,20]. In this study, we investigated the cell wall composition and dynamics of *C. vulgaris* in three different conditions: in exponential phase, stationary phase, and salinity stress condition (0.1 M NaCl). For that, the approach that we developed combined three types of analysis. First, AFM was used to image the cells and probe the cell wall roughness and nanomechanical properties. Then, XPS analysis was used to give a global view of the cell wall composition and determine this way the relative amounts of the three fractions, proteic, saccharidic, and lipidic. Finally, to give a complete view of the cell wall composition, chemical hydrolysis followed by HPLC was performed to determine the saccharidic composition of the cell wall. In the end, the combination of these three techniques allow to get a complete picture of the effects of culture condition on the cell wall composition and dynamics of *C. vulgaris* cell wall, but also to understand the link between composition and architecture and the effects of composition changes on cell surface biophysical properties. These results provide important information that can be further used to develop more efficient and targeted lipid extraction methods for industrial applications, but also to better apprehend the microalgae cell wall and its interaction with its environment.

## MATERIALS AND METHODS

### Microalgae strain and culture

The green freshwater microalgae *Chlorella vulgaris* strain CCAP 211/11B (Culture Collection of Algae and Protozoa, Scotland, UK) was cultivated in sterile conditions in Wright’s cryptophyte (WC) medium prepared with deionized water, as previously described [21]. Cells were cultivated at 20°C, under 120 rpm agitation, in an incubator equipped with white neon light tubes providing illumination of approximately 40 μmol photons m^-2^ s^-1,^ with a photoperiod of 18h light: 6h dark. Exponential phase experiments were carried out with 7 days batch cultures, whereas stationary phase and salinity stress condition (0.1M NaCl) experiments were carried out with 21 days batch cultures. Cell growth was monitored (cell abundance) for three different cultures in each condition.

### AFM imaging

Before experiments, cells were first harvested by centrifugation (3000 g, 3 min), washed two times in PBS at pH 7.4, and immobilized on polyethylenimine (PEI, Sigma-Aldrich P3143) coated glass slides prepared as previously described [22]. AFM images of *C. vulgaris* cells were then recorded in PBS at pH 7.4, using the Quantitative Imaging mode available on the Nanowizard III AFM (Bruker, USA), with MSCT cantilevers (Bruker, nominal spring constant of 0.01 N/m). Images were recorded with a resolution of 150 pixels × 150 pixels, at an applied force of <1.0 nN and a constant approach/retract speed of 90 μm/s (z-range of 3 μm). In all cases the cantilevers spring constants were determined by the thermal noise method prior to imaging [23]. In each case, 3 cells were imaged.

### Roughness analyses

Roughness analyses were performed on *C. vulgaris* cells immobilized on positively charged glass slides (Superfrost™ Plus adhesion, Epredia, USA). High resolution images of the cell walls were recorded in PBS using QI advanced imaging mode available on the Nanowizard III AFM (Bruker, USA), using MSCT cantilevers (Bruker, nominal spring constant of 0.01 N/m). In each case, 13 cells coming from at least 2 independent culture were imaged and images were recorded with a resolution of 150 x 150 pixels using an applied force < 1 nN. In all cases the cantilevers spring constants were determined by the thermal noise method prior to imaging [23]. The height images obtained were then analyzed using the Data Processing software (Bruker, USA) to determine the arithmetic average roughness (Ra)..

### Nanomechanical Analyses

For nanoindentation experiments, the AFM was used in force spectroscopy mode using an applied force comprised between 0.5 and 2 nN depending on the condition, with MSCT cantilevers (Bruker, nominal spring constant of 0.1 N/m). In each case, 12 cells coming from at least 3 independent culture were analyzed (approximately 600 force curve for each cells, details are given in the Results and Discussion section). Young’s moduli were then calculated from the indentation curves obtained (50 nm long indentation segments were used) using the Hertz model in which the force F, indentation (δ), and Young’s modulus (Ym) follow equation 1, where α is the tip opening angle (17.5°), and ν the Poisson ratio (arbitrarily assumed to be 0.5). The cantilevers spring constants were determined by the thermal noise method [23].

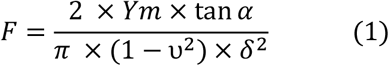

### Isolation of microalgae cell walls

The isolation process is based on the study by Schiavone *et al.* [24]. Briefly, cells coming from at least 3 independent culture in each case were harvested by centrifugation (4700 g, 10 min) and washed 2 times with sterile deionized water. Then the pellets were resuspended in 10 mL of sterile water and transferred to a 15 mL falcon tube containing 2 g of acid washed glass beads (0.5 mm of diameter, Thermofisher, G8772-100G). Cells were then disrupted using a Fastprep system (MP Biomedicals): 10 cycles of 20 s with intervals of 1 min were performed while keeping the pellets on ice. The cell suspension was directly collected, and the glass beads in the pellet were extensively washed with cold deionized water. The supernatant and washings were pooled and centrifuged at 4700 g for 15 min at 4 °C. The cell wall-containing pellet was again washed two times with cold deionized water. Then, pellets were frozen at −80°C and lyophilized in a freeze dryer until complete dryness.

### Acid hydrolysis of microalgae cell walls and quantification of carbohydrates by HPAEC-PAD

Sulphuric acid (72 %) hydrolysis of the cell wall was carried out as described previously [24,25]. Briefly the freeze-dried biomass (10 mg) was suspended in 75 μL of 72% H_2_SO_4_ and vortexed to dissolved the powder. After 3 hours of incubation of the suspension at room temperature (20-25 °C) (vortex every 30 min) sample were diluted to reach 2N H_2_SO_4_ and incubated in a sand bath for 4 hours at 100°C (vortex every 1h). This was followed by a neutralization step with 40 g/L Ba(OH)_2_, then, samples were filled with water until 25 mL volume is reached. Finally the suspensions were centrifuged (10 min 3000 g) and supernatants were collected. The supernatant was filtered on a 0.2 μm Amicon and then was analyzed by high-performance anion exchange chromatography (HPAEC) on ICS 3000 system (Thermofisher Scientific, France). Separation of the released monosaccharides (glucose, galactose, rhamnose, arabinose, glucosamine, xylose, mannose) were performed on a CarboPac PA1 analytical column (250 x 4 mm) with a guard column CarboPac PA1 using an isocratic elution at 18 mM NaOH (200 mM) for 20 min at 1 mL/min and 25°C. After a washing step was performed with 200 mM NaOH for 5 min and 300 mM sodium acetate in 200 mM NaOH for 10 min, following a cycle of equilibration of the column with 18 mM NaOH for 20 min. Sugar residues were detected on a pulsed amperometric system equipped with a gold electrode and a reference electrode (Ag/AgCl) using the method “Carbohydrate standard quadruple potential”.

### XPS analysis

The photoelectron emission spectra were recorded using a monochromatised Al Kalpha (hv = 1486.6 eV) source on a ThermoScientific K-Alpha system. The X-ray Spot size was about 400 μm. The Pass energy was fixed at 30 eV with a step of 0.1 eV for core levels and 160 eV for surveys (step 1 eV) The spectrometer energy calibration was done using the Au 4f7/2 (83.9 ± 0.1 eV) and Cu 2p3/2 (932.8 ± 0.1 eV) photoelectron lines. XPS spectra were recorded in direct mode N (Ec) and the background signal was removed using the Shirley method. The flood Gun was used to neutralize charge effects on the top surface. For each condition, experiments were performed in triplicate on cell walls isolated from cells coming from at least 2 independent cultures were conducted.

### Protein quantification

Protein quantifications were performed with cells coming from at least 3 independent cultures in each case using a bicinchoninic acid (BCA) protein assay kit (ThermoScientific, 23225) according to Smith *et al.* [26]. Bovine serum albumin (BSA) was used as a standard and the protocol was followed at 60 °C for 30 minutes following the manufacturer guidelines.

### Hydrophobicity measurements

Hydrophobicity measurements were conducted using AFM and FluidFM as described in Demir *et al.* [27] Briefly, an air-bubble was produced using a Nanowizard III AFM (Bruker, USA), equipped with FluidFM technology (Cytosurge AG, Switzerland). Experiments were performed in PBS, using microfluidic micropipette probes with an aperture of 8 μm (spring constant of 0.3, and 2 N/m, Cytosurge AG, Switzerland). The probes were calibrated using the thermal noise method prior to measurement.[23] *C. vulgaris* cells were first harvested by centrifugation (3000 g, 3 min), washed two times in PBS at pH 7.4 and immobilized on positively charged glass slides (Superfrost™ Plus adhesion, Epredia, USA). Interactions between the formed bubbles at the aperture of the microfluidic micropipette probes and 8 cells coming from at least 2 independent culture were measured in force spectroscopy mode using a constant applied force of 1 nN. Force curves (approximately 625 force curve for each cell) were recorded with a retraction z-length of up to 3 μm and a constant retraction speed of 3.0-6.0 μm/s. The adhesion force between bubble and *C. vulgaris* cell wall corresponds to the height of the adhesion peak.

### Statistical analysis

Experimental results represent the mean ± standard deviation (SD) of at least three replicates. For each experiments, the number of replicates is indicated both in the Material and Methods section in the corresponding paragraphs, and in the Results and Discussion section. For large samples (n>20 values) unpaired student t-test was used to evaluate if the differences between the conditions are significant. For small samples (n<20 values) non-parametric Mann and Whitney test was used to assess the differences. The differences were considered significant at p <0.05.

## RESULTS AND DISCUSSION

### Influence of salinity stress on growth of *C. vulgaris*

A first step in the study was to evaluate the effects of the culture conditions used on cell growth over time. For that, *C. vulgaris* cells were cultured during 30 days in normal conditions or under salinity stress; the optical density (OD) of the suspensions was measured every day to monitor cell growth. The results are presented in Figure 1. A first information from these growth curves is that both in control and in salinity stress condition, they do not show a lag phase, meaning that cells did not need time to adapt to the presence of salt in the medium. *C. vulgaris* cells cultured in standard conditions were in an exponential growth phase during 21 days, before reaching the stationary phase. However, when cells were cultivated under salinity stress, the exponential growth phase was significantly reduced to 15 days. Then cells stayed in stationary phase during 10 more days, after that cell concentration decreased by 15%, indicating partial cell death. This decline phase is not observed for cells cultivated in standard conditions. Studies in the literature have determined that the NaCl present in the medium, at a certain concentration, becomes toxic and reduces the growth, explaining the decline in saline stress condition. For instance Singh and coworkers reported similar growth curves patterns with a decline for *C. vulgaris* cells cultivated in saline stress conditions using, different salt concentrations [28]. These measurements thus show that culture conditions have an important impact on cell growth, and thus most likely on cell wall structure and composition. In addition, it has been showed that cells in stationary phase undergo a pH increase that modifies cell wall properties, such as composition [29] and architecture [22,30]. Thus in the next part of this work, we used AFM, XPS and chemical hydrolysis followed by HPLC to analyze the cell wall of cells grown in standard conditions in both exponential (7 days of culture) and stationary phase (21 days of culture), and of cells grown in saline stress (0.1 M of NaCl) during 21 days. Analysis of *C. vulgaris* cultivated with 0.1M NaCl (salinity stress condition) were performed at 21 days with stationary phase cells, where the highest difference in growth is observed compared to cells in cultured in normal conditions.

**Figure 1.**
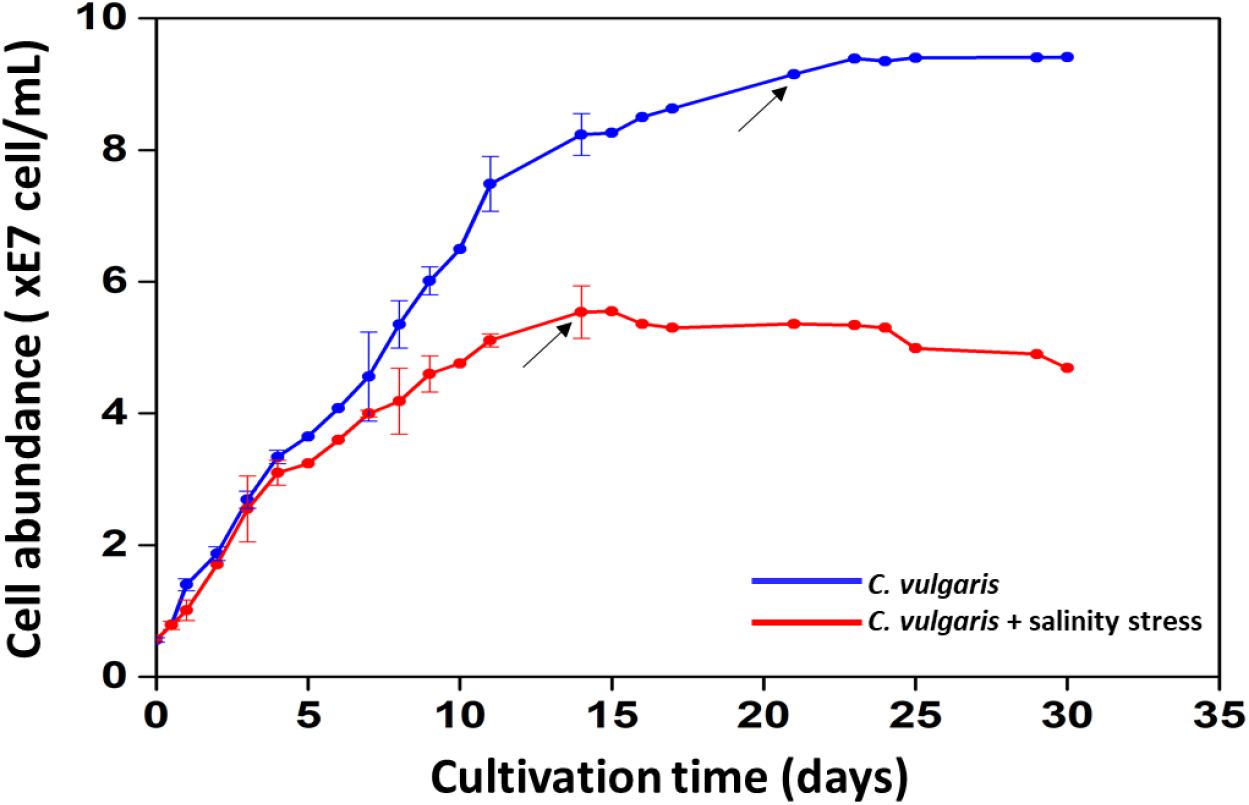
C. vulgaris growth. Variation in cell abundance (cell/mL) measured over time for batch cultures of C. vulgaris in standard conditions (WC culture medium, blue curve) and in salinity stress condition (WC culture medium supplemented with 0.1 M NaCl, red curve). The black arrows indicate in each case the end of exponential phase and the beginning of stationary phase.

### Probing the biophysical properties of cell surfaces in the different culture conditions using AFM

Then in a next step, *C. vulgaris* cells grown in the different conditions described above were analyzed using AFM. For that, first, height images of the whole cells were obtained using a force spectroscopy-based imaging mode (Quantitative Imaging mode, QI [31]) with a resolution of 150 x 150 pixels. Images are shown in Figure 2a, b and c for exponential phase, stationary phase and salinity stress condition respectively. For cells grown in standard conditions, no significant morphological changes could be observed between the two different growth stages (exponential phase and stationary phase). However, under saline stress, defects at the cell surface can be observed; the cell wall appears to rougher compared to cells grown in standard conditions. To quantify this, we then made zoom-in high resolution images on small areas on top of cells (300 nm x 300 nm) using QI imaging mode, as shown in Figure 2d, e and f for exponential phase, stationary phase and salinity stress condition respectively. The cross-sections (Figure 2g-i) taken along the white lines on these images show that surface morphology is modified both for stationary phase cells and saline stress cells compared to exponential phase cells. In the case of the salinity stress, even larger patterns are visible on the cross-section indicating a larger deformation of the cell surface. To quantify these deformations, the average roughness Ra of the surface was measured directly from the height images presented in Figure 2d-f. In each condition, roughness measurements were performed on 13 different *C. vulgaris* cells coming from at least two independent culture; the results are shown in the box plot in Figure 2j. This quantitative analysis shows that in exponential phase, cells have an average roughness of 1.1 ± 0.4 nm which increases to 1.5 ± 0.7 nm in stationary phase and to 1.7 ± 1.2 nm in salinity stress condition. A statistical analysis (Mann-Whitney test) shows that these differences are not significant. This is in line with the existing literature; for instance, similar roughness values were also recorded for *Dunaliella tertiolecta* cells grown in exponential and stationary phase [32]. These results prove that the growth stage of cells does not have a significant effect on the cell surface roughness. However, when we compare the distribution of the roughness values measured in each case, in salinity stress condition the standard deviation is higher (1.2 nm) compared to stationary phase (0.7 nm) or exponential phase cells (0.4 nm). Thus even though the differences between conditions are not significant, still, applying a stress seem to increase the heterogeneity of the surface roughness values measured on different cells. A modification of cell surface roughness for cells under stress condition has already been showed for other types of microorganisms. For example, Schiavone *et al.* studied the effects of ethanol stress on the yeast *Saccharomyces cerevisiae* and reported a 50% increase in cell surface roughness when cells were submitted to this stress [33]. In our case, the increase in the heterogeneity of the roughness values that we observe in salinity stress condition could indicate that more molecules protrude from the cell surface, as it was hypothesized in a previous AFM study on *C. vulgaris* [21] or that charged surface molecules get coiled because of the salt present in the medium which could also result in a change in surface roughness [34].

**Figure 2.**
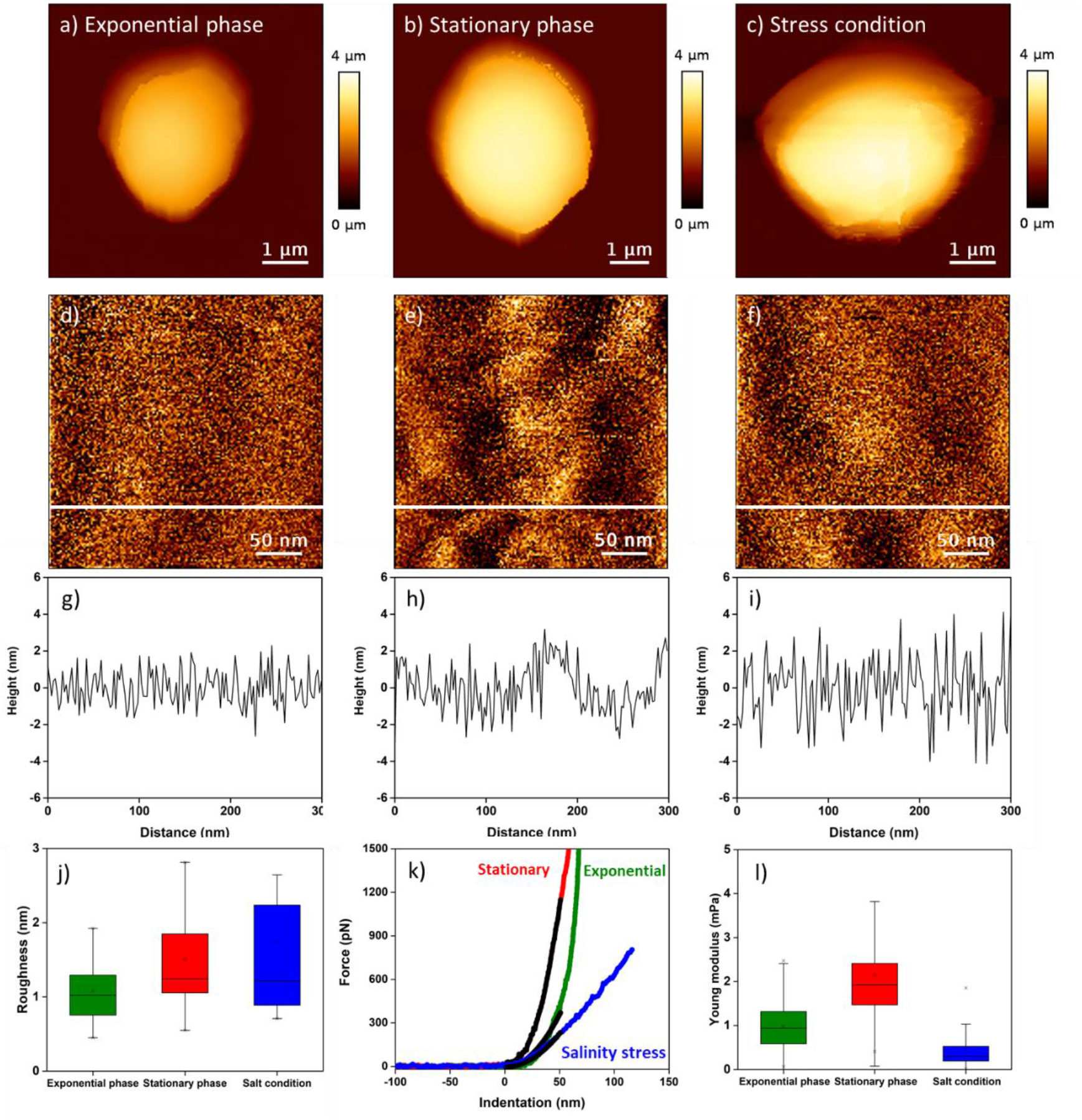
Roughness and nanomechanics of C. vulgaris cell wall. AFM images of single C. vulgaris cell in a) exponential phase b) stationary phase and c) salinity stress condition. AFM height images recorded on an area of 300 nm x 300 nm on top of cells in d) exponential phase e) stationary phase and f) salinity stress condition. g), h) and i) are cross-section taken along the white lines in d), e) and f) respectively. j) is a box plot showing the distribution of C. vulgaris surface roughness in exponential phase (green box), stationary phase (red box) and salinity stress conditions (blue box). k) Indentation curves (green, red and blue lines) fitted with the Hertz model on a 50 nm indentation segment (black lines) recorded on top of C. vulgaris cells in exponential phase (green curve), stationary phase (red curve) and salinity stress conditions (blue curve). l) Boxplot showing the distribution of Young’s modulus values measured on top of C. vulgaris cells in exponential phase (green box), stationary phase (red box) and salinity stress conditions (blue box).

Other properties that we can measure using AFM to get information on the structure or architecture of the cell wall are the nanomechanical properties. To obtain quantitative information on the nanomechanical properties of *C. vulgaris* cell wall, we determined the Young’s modulus (Ym) using nanoindentation measurements (Figure 2 k and l). In this type of measurement, a cantilever with known mechanical properties, is pressed against the cell surface at a specific force. This allows extracting the Ym of the cell wall, in other words its compression resistance. The Ym is thus a value that reflects the cell wall rigidly; the higher the Ym value, the more rigid the cell wall. In this study, nanoindentation measurements, which provide access to force *versus* distance curves, were performed on areas of 300 nm × 300 nm on top of cells, on 12 cells coming from at least three independent cultures. Ym values were then obtained first by converting the force curves into force *versus* indentation curves as shown in Figure 2k, and then by fitting them with a theoretical model, in our case, the Hertz model [35] (black circles on the curves in Figure 2k). Nanoindentation curves, obtained on cells in the different conditions, show a different slope, meaning that the AFM probe does not indent the same way in each case. For instance, it is able to indent deeper in exponential phase cells compared to stationary phase, meaning that this change in the growth phase increases the rigidity of the cell wall. The indentation is even deeper for cells cultured in salinity stress condition (21-days of culture) compared to stationary phase cells, showing that addition of salts have a direct impact on cell wall rigidity. Quantitative analysis of the Ym extracted from thousands of force curves recorded on 12 cells in each condition confirm these findings and show that exponential phase cells have an average Ym of 981.6 ± 554.5 kPa (n = 6011 force curves), which increases to 2.1 ± 1.3 MPa for stationary phase cells (n = 6580 force curves, the difference is significant at a p-value of 0.05, unpaired t-test). These values are comparable with previous nanomechanical measurements performed on *C. vulgaris* [21]. Moreover, a difference in the rigidity of cells in the two growth phases has already been observed for another microalgae species: for instance Pillet and co-workers investigated the nanomechanical properties of *D. tertiolecta* in exponential and stationary phase and found that cells in stationary phase are softer [32]. These modifications of rigidity over time can be explained by the fact that the pH of microalgae cultures changes with time. Indeed, several studies have showed that pH changes have a direct effect on microalgae cell wall nanomechanical properties [21,30]. For instance in our case, in normal conditions, the initial pH of *C. vulgaris* cultures is of 7.8, and then increases to 9.0 at the end of exponential phase (7 days) and to 8.5 in stationary phase (21 days). For cells submitted to saline stress, the Ym value this time drops to 433.2 ± 415.9 kPa (n = 7005 force curves, the difference is significant at a p-value of 0.05, unpaired t-test). This difference in the Ym value in this case can be explained by the direct impact of the environmental stress on the cell wall, as other studies have showed. For example, Yap *et al.* found that *Chlorococcum sp.* cells submitted to a N-deprivation had a Ym approximately 30% higher than for N-replete cells [36]. However, it could also be due to the osmotic pressure; in this saline condition, water may flow out of the cell, thereby changing its turgor pressure and thus the Ym of the cell wall [37]. To verify this point, we measured the diameter of different cells in all three conditions; no significant difference in the diameters measured were observed (Supplementary Figure S1). To understand if the changes observed in the roughness or in the rigidity of cells in stationary phase can be linked to the cell wall architecture, we need to determine the biochemical cell wall composition. Such correlation between cell surface biophysical properties and cell wall composition has been made already for different microorganisms such as yeasts [15], and might also be true in the case of this study.

### Biochemical composition of microalgae *C. vulgaris* cells based on XPS

Thus to explore the biochemical composition of *C. vulgaris* cell wall depending on the different culture conditions used in this study, we used XPS. XPS technique quantitatively measures the elemental composition of a surface, biotic or abiotic, including the chemical functionalities in which the elements are involved [38]. XPS has proven to be a powerful technique to determine the cell wall composition of yeast and bacteria; in the case of microalgae, the few available studies use cryo-XPS, whose difference with conventional XPS relies in the sample preparation procedure.^18^ In the case of our study, the XPS analysis were performed on isolated freeze-dried cell walls. Because of XPS probing depth of less than 10 nm, and the thickness of *C. vulgaris* cell wall being of approximately 60 nm [6], using whole cells would provide information only on the near-surface region of cell wall. This is why we chose to work with isolated cell walls: by performing several measurements for each conditions we can this way have a more global information on the composition of the whole cell wall as they are mixed and not only its surface. On XPS spectra, the position of the XPS peak is known to be dependent on the chemical environment of the element, the binding energy having a tendency to decrease as the electron density on the atom increases [39]. XPS overall spectra to identify the elements are shown in supplementary Figure S2. Moreover, carbon, nitrogen and oxygen spectra and elemental atomic percentages obtained for cells in exponential phase, stationary phase and salinity stress conditions are presented in Supplementary Figures S3, S4 and S5 as well as Tables S1, S2 and S3 respectively. Figure 3 presents the XPS carbon 1s spectra recorded on *C. vulgaris* cells in exponential phase, in stationary phase and in salinity stress condition. The positions, relative intensities and average atomic percentages (out of three replicates) of the carbon peaks are presented in the spectra.

**Figure 3.**
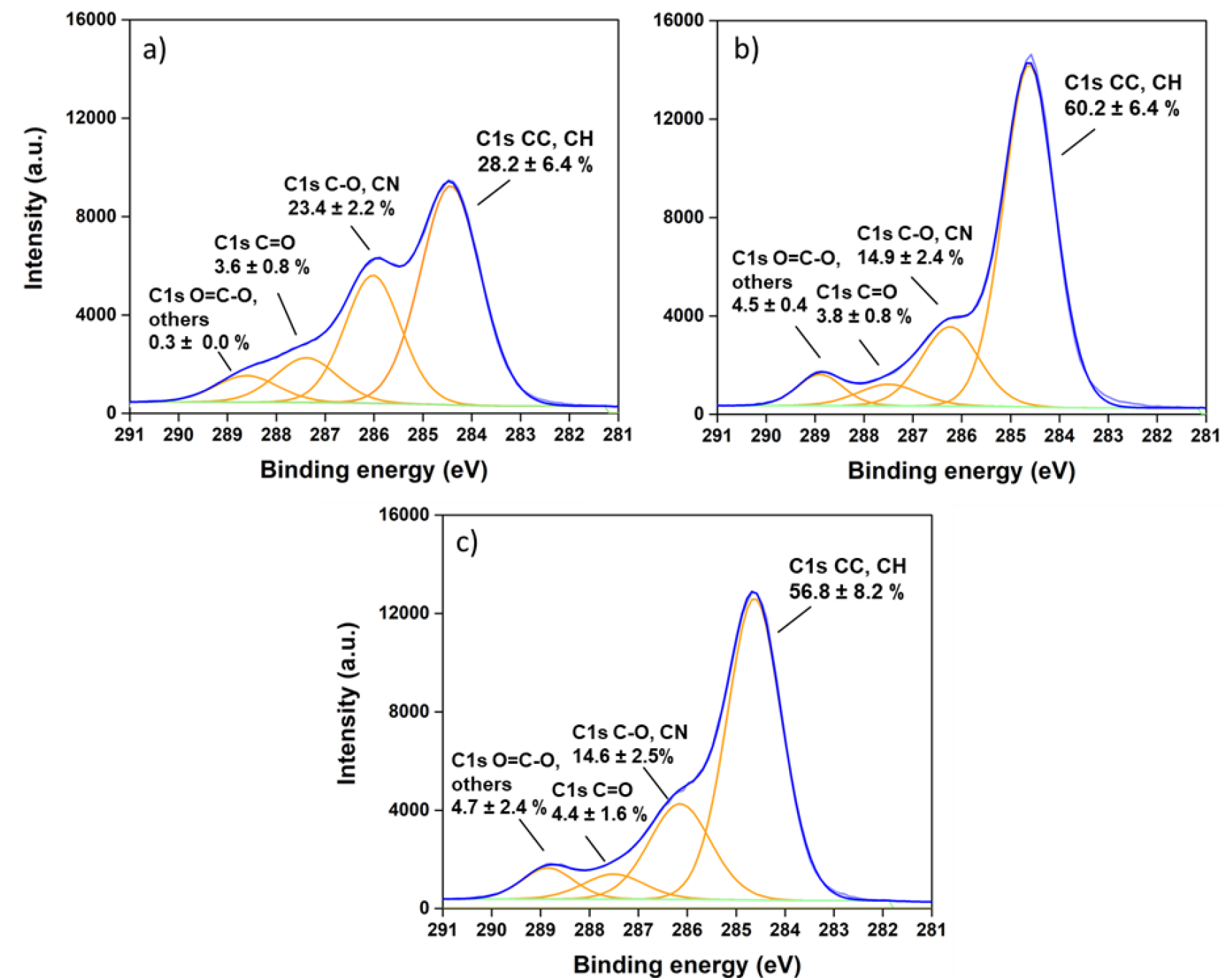
XPS analysis of C. vulgaris cell wall. Carbon 1s peaks recorded on C. vulgaris cell walls isolated from cells grown in standard condition in a) exponential phase, b) stationary phase and c) salinity stress condition. Average atomic percentages and standard deviations were calculated from triplicates (n=3)

In each case, one spectra representative of all the measurements performed is presented; the mean atomic percentages obtained for the three different measurements performed in each conditions are indicated. Exponential phase, stationary phase and salinity stress condition showed reproducible surface composition between the different cultures; the standard deviations obtained for the atomic percentages reflect the normal heterogeneity found between different biological cultures. The average atomic percentage of C–C components increases by a factor of 2 from exponential phase (28.2 ± 6.4) to stationary phase (60.2 ± 6.4), and then slightly decreases from stationary phase to salinity stress condition (56.8 ± 8.2). Regarding C-O components, their average atomic percentage decreases by a factor of 1.6 from exponential phase (23.4 ± 2.2) to both stationary phase (14.9 ± 2.4) and salinity stress condition (14.6 ± 2.5). C=O components stay relatively constant between the three conditions, while the atomic percentage of O-C=O components is 15 times higher in exponential phase (0.3 ± 0.0) compared to stationary phase (4.5 ± 0.4), and slightly further increases when cells are exposed to salinity stress condition (4.7 ± 2.4). To understand the implications of these changes in terms of cell wall composition, we analyzed the XPS data using theoretical models developed by Rouxhet and coworkers [39]. For many biological systems, including the microalgae cell wall, three main classes of model compounds can be considered: proteins (Pr), polysaccharides (Po), and lipids (HC). The authors proposed a set of equations that allows to evaluate the proportion of carbon associated with these three model compounds, based on the three main components of the carbon peak. This model predicts that:

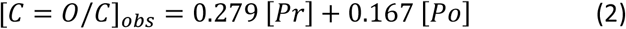

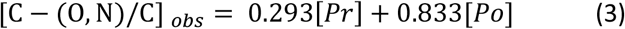

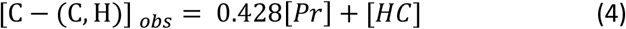

Simultaneously solving these carbon base equations gives the proportion of proteins, polysaccharides and lipids present in the cell wall of *C. vulgaris* in exponential phase, stationary phase and salinity stress condition; the results obtained are presented in Figure 4 and summarized in Supplementary Table 4.

**Figure 4.**
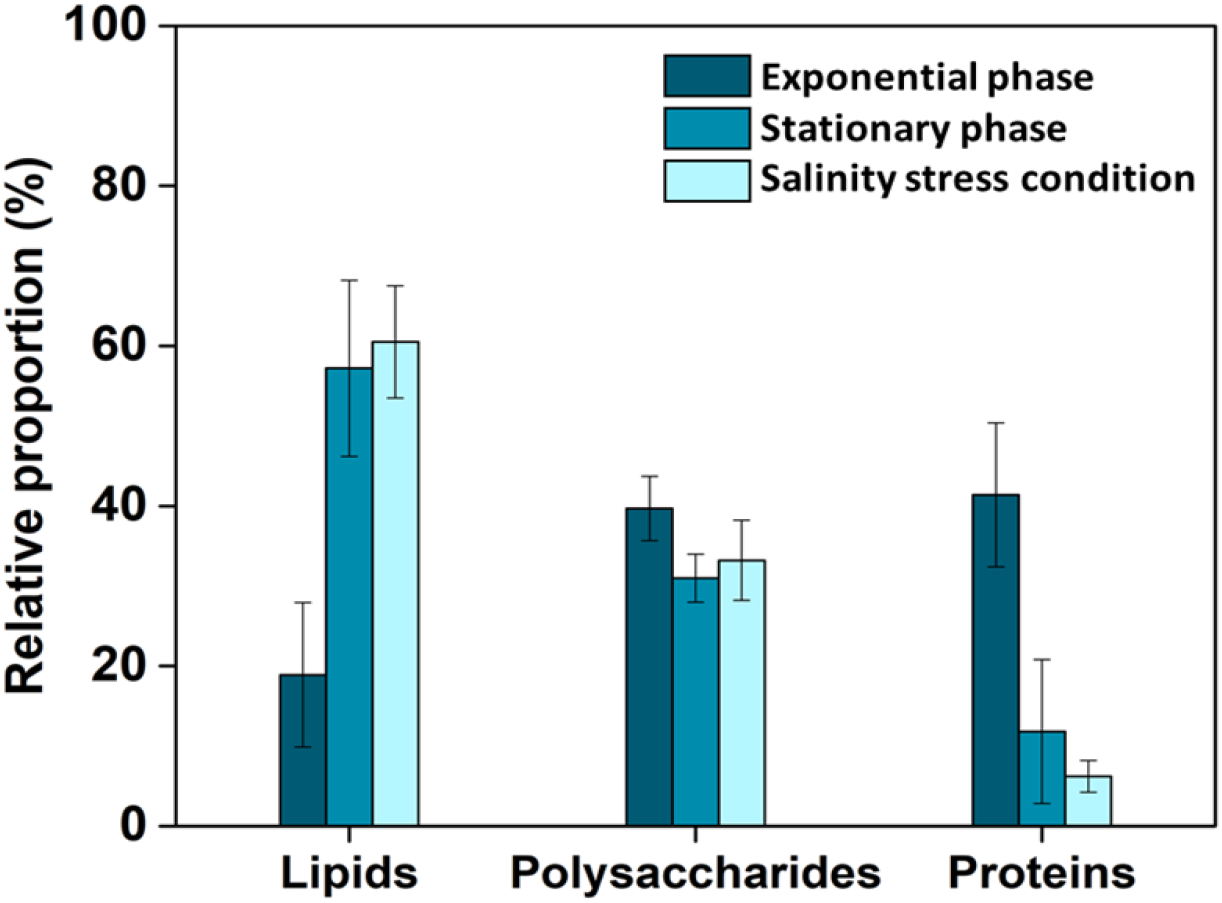
Biochemical composition of C. vulgaris cell wall. This histogram shows the relative proportions of carbon associated with lipids, polysaccharides and proteins in the cell wall of cells in exponential phase, stationary phase and salinity stress condition (0.1M NaCl). The XPS values are here normalized to 100% using a calculation based on the carbon concentration in each type of constituent (mmol of carbon/g of constituent) [39] and represent the relative proportion of the corresponding compounds. The error bars indicate the deviation from the average of the triplicates (n=3).

In exponential phase, the dominant constituents in the cell wall of *C. vulgaris* are proteins (41.4 ± 9.2%) and polysaccharides (39.7 ± 3.5%). This is in line with the study by Shchukarev *et al.*, where the authors also found that these compounds were predominant in the cell wall of *C. vulgaris* using cryo-XPS [13]. With aging from exponential phase to stationary phase, the difference in the cell wall composition is pronounced. The lipid content increases by a 3.0-fold at the expanse of a 3.5-fold decrease in the protein content. The polysaccharide proportion decreases as well but in a non-significant manner. This increase in the lipid content is even more pronounced with cells exposed to the saline stress, in this case also at the expanse of a decrease in the protein content. The proportion of polysaccharides, as for it, remains approximately the same compared to stationary phase cells. To make sure these analysis are correct, we also determined the protein concentration in the cell walls from cells in the three conditions using an assay kit. The results obtained (Supplementary Figure S6) showed that the highest protein concentration is observed for exponential phase cells and decreases importantly, in the same proportions than observed using XPS, for stationary phase and salinity stress condition. This additional experiment thus gives confidence in the validity of the XPS measurements performed.

These results are quite interesting. Indeed, it has been showed by multiple studies that salt stress induces the accumulation of lipids in *C. vulgaris*, more specifically of storage neutral lipids (TAG) [40,41]. These neutral lipids are present in the cells as droplets in the chloroplast matrix and in the cytoplasm [42]. Thus by definition, they should not be present in the cell wall. The lipids that have a structural role and that are located on the cell wall are polar lipids, which are mainly phospholipids and glycolipids. But while the effects of salinity stress on storage lipid accumulation has been investigated before, its effects on the production of polar lipids composing the cell wall has never been studied, as far as we know. The XPS data obtained here seem to indicate that their production is also increased in stationary phase or under salinity stress. To prove this, a way is to evaluate the hydrophobic properties of the cell wall, as lipids are the only components that can provide hydrophobic properties to cells. Indeed, polysaccharides are hydrophilic, and while proteins could also have hydrophobic properties, their relative fraction in the cell wall in stationary phase and in salinity stress condition is small, making it unlikely that they could participate in a significant manner to the hydrophobicity of the cell wall. Hydrophobic organic material are mostly composed of large fractions of aliphatic or aromatic substances. Thus the ratio of aliphatic carbon components to the total carbon in the C1 spectra can be directly linked to the hydrophobicity of the surface [39,43,13]. This method to determine hydrophobicity has already been applied in different studies, for example, to compare the hydrophobicity of bacteria between aqueous phase and organic phase [43], or to compare the relative hydrophobicities of different microalgae species [13]. In the case of our study, the aliphatic content in *C. vulgaris* in exponential phase corresponds to 42.9 ± 7.0 % of the total carbon, which is lower than for the other two conditions, where the ratios are of 71.9 ± 5.1% in stationary phase and of 70.3 ± 7.2 % in salinity stress condition. This means that cells are more hydrophobic in stationary phase and salinity stress condition compared to exponential phase, most likely because of the increased amount of lipids present in the cell wall in these conditions.

To confirm the XPS data and further prove that the amount of structural lipids in the cell wall is increased in stationary phase and salinity stress conditions, we performed another type of experiments to evidence their presence in the cell wall. For that, we probed the hydrophobic properties of the surface of living cells in the different conditions using a method recently developed in our team based on FluidFM [44], which combines AFM and microfluidics. It consists in producing a bubble at the aperture of a FluidFM cantilever, and probing its interactions with cells in force spectroscopy experiments [27]. As bubbles in water behave like hydrophobic surfaces, the interactions recorded directly reflect the hydrophobic properties of cells. The higher the adhesion force, the more hydrophobic the surface is. The results are presented in Figure 5, a schematic representation of the experiment’s principle is shown in Figure 5a. For cells in exponential phase, force curves show a single retract peak at the contact point, typical of hydrophobic interactions [45], (inset in Figure 5b) with an average force of 3.7 ± 0.7 nN (Figure 5b, n= 4950 force curves obtained from 8 cells coming from 2 independent cultures). The same type of adhesion peak is observed in stationary phase and in saline stress conditions, but in these cases, the adhesion force is increased to 4.9 ± 0.8 nN for cells in exponential phase (Figure 5c, n= 4938 force curves obtained from 8 cells coming from 2 independent cultures), and to 5.6 ± 1.1 nN in saline stress conditions. (Figure 5d, n= 4774 force curves obtained from 8 cells coming from 2 independent cultures). These values are all significantly different at a p-value < 0.05 (unpaired student test). These results are important; they were performed on live cells cultivated in the different conditions used in this study, and also show that cells are more hydrophobic and thus that more lipids are present in stationary phase and in saline stress conditions. Thus they confirm the XPS data obtained and indeed, in these conditions, not only the production of storage lipids by the cells is increased, but also the production of structural lipids present in the cell wall.

**Figure 5.**
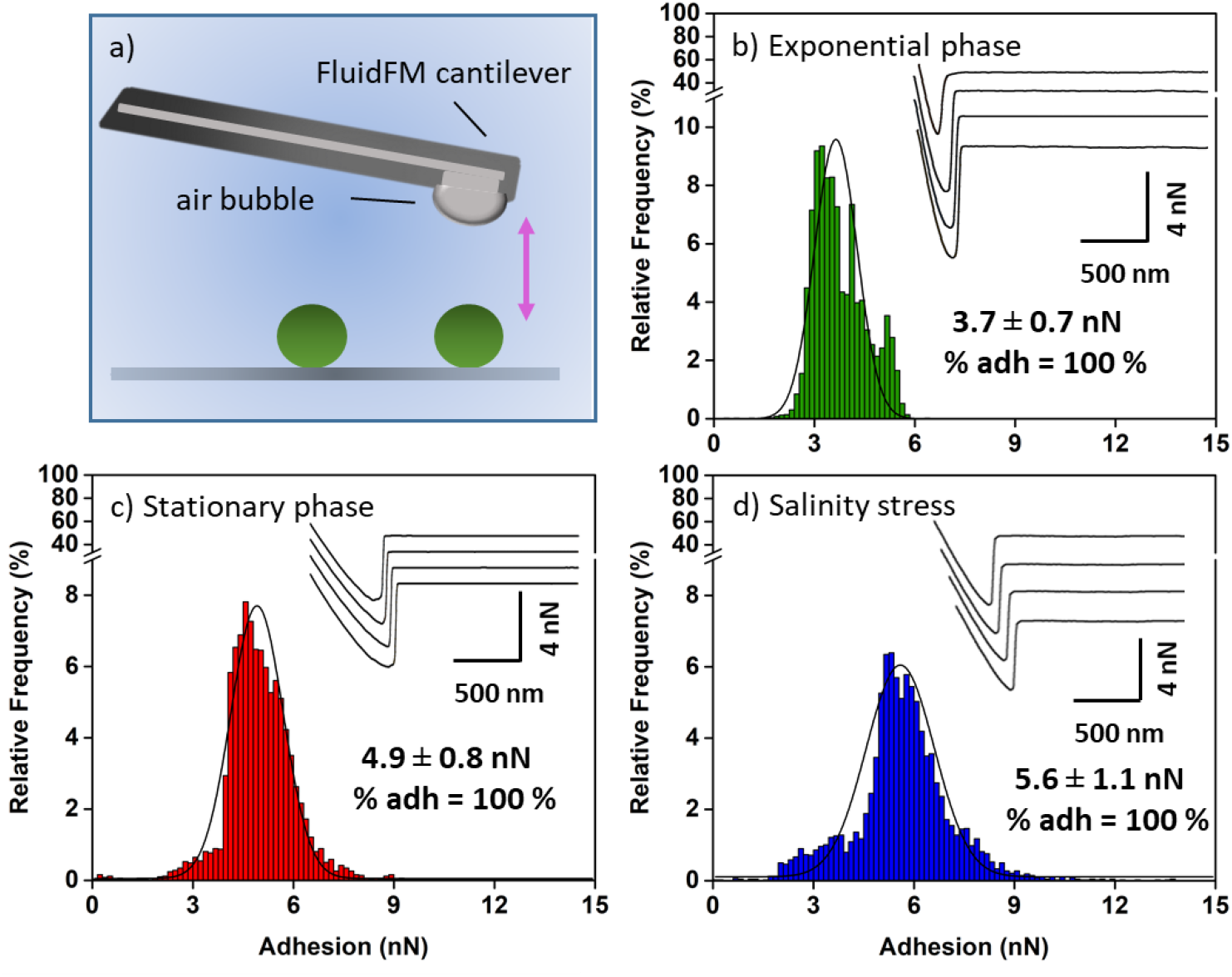
Probing the hydrophobic properties of C. vulgaris cell wall. a) Schematic representation of the experiment’s principle: the interactions between bubbles produced at the aperture of FluidFM cantilevers and C. vulgaris cells immobilized on a surface are probed in force spectroscopy mode. Adhesion force histogram obtained between bubble and C. vulgaris cells in b) exponential phase c) stationary phase and d) salinity stress condition. Insets in b), c) and d) show representative force curves obtained in each case.

Regarding the variations observed in the proteic content in the cell wall, the decrease of this fraction can be explained by the fact that in stationary phase or under salt stress condition, the photosynthesis is inhibited [46]. This has for consequence to decrease the chlorophyll content in cells which results in cell growth inhibition, as we could observe on the growth curves (Figure 1), but also in the decrease of the proteic content of cells. This has already been showed for *C. vulgaris* cells submitted to the same saline stress (0.1 M) as in this study [28]. In addition, a genomic study has showed that under saline stress conditions, the expression of a large number of genes was down-regulated, in particular genes involved in the photosystem light-harvesting pathways as well as genes involved in protein synthesis and stability [47].

Altogether collecting these information on the cell wall composition allows understanding better the biophysical observations made with AFM in terms of cell wall roughness and rigidity. For instance, XPS results show that the proportion of polysaccharides does not vary much; polysaccharides are long polymers that can be exposed at the outer surface of cells, and thus which can be responsible for the cell wall roughness [48,49]. Our AFM data showed no significant difference in the roughness between the three conditions; this could perhaps be due to the fact that the proportion of this fraction remains similar in the different conditions tested. Also the rigidity of cells in stationary phase is significantly more important than for cells in exponential phase; in this case the change in the rigidity can be attributed to the different cell wall composition between these two conditions. This is an interesting point because it shows that the cell wall is a dynamic and complex structure able to rearrange during the growth of cells. Cells in stationary phase and in saline conditions have a similar cell wall composition, but a very different rigidity. In saline condition, the decrease in the rigidity of the cells can probably be mainly attributed to a decrease of turgor pressure. However, another hypothesis could also be that in this case, because of the environmental stress applied, the cell wall rearranges with a different architecture than for cells in stationary phase. This different architecture, for a same composition, could also contribute to the variation in the rigidity observed. Thus taken together, the combination of XPS data with AFM analysis of the cell wall enlightens its complexity and dynamics. These are important information for example to optimize disruption procedures to extract the lipid content of cells. For instance, if mechanical disruption is used, using a saline stress on cells to produce lipids might be a better alternative; cells are less rigid in this condition and can be more easily ruptured. In addition, different disruption procedures using enzymatic degradations, or a combination of enzymatic degradation and mechanical rupturing can be used; to optimize such procedures and select adapted enzymes, more information is needed on the polysaccharides present in the cell wall.

### The saccharidic composition of *C. vulgaris* cell wall is influenced by the growth phase and culture conditions

Thus to determine the polysaccharidic composition of the cell wall of *C. vulgaris* in the different conditions, isolated cell walls were used for acid hydrolysis using the well-known concentrated sulfuric acid method. The monosaccharide composition of microalgae cell walls is reported in Figure 6 (HPAEC-PAD spectra are presented in Supplementary Figure S3); in this figure the monomer concentrations are expressed as (w/w) mg of monomer per gram of dry cell wall (DCW). Our results show that *C. vulgaris* cell wall is composed predominantly of glucose followed by galactose, rhamnose, arabinose and glucosamine. Other monosaccharides are also present but in smaller amounts; xylose and mannose, although for those, the peaks on the spectra overlap (see HPAEC-PAD spectra in Supplementary Figure S3), meaning that their quantification is not feasible. Glucose and galactose represent the main components of *C. vulgaris* carbohydrates biomass in all three conditions, and account for 78% of the total amount of cell wall carbohydrates in stationary phase and in salinity stress conditions, and for 86% in exponential phase. Therefore, the main composition changes taking place in the different conditions thus concern these two sugars. Indeed, glucose concentration drops from 96.0 mg/g in exponential phase to 55.3 mg/g in stationary phase and 40.1 mg/g in saline stress condition. While the relative quantities are less important for galactose, the same tendency is observed, the concentration decreases from 18.4 mg/g for exponential phase cells to 12.8 mg/g and 9.8 mg/g for stationary phase and salinity stress conditions cells respectively. The concentrations of the other monosaccharides present, rhamnose, arabinose and glucosamine remain almost the same in the different conditions. Regarding xylose and mannose, while no concentration values can be given in this case, the comparison of the total peak area (combination of xylose and mannose) shows an increase of almost 80% in the total concentration of these sugar from exponential to stationary phase and then shows a decrease of almost 50% from stationary phase to salinity stress condition. As already mentioned the protein concentration was determined using the bicinchoninic acid method (Supplementary Figure S6), thus the proportion of lipids can be deduced. These proportions obtained based on these results are presented in Supplementary Figure S8; they follow the same pattern than what was obtained using XPS data, thus confirming the validity of our observations. Note that the percentages obtained for each fractions are different in the two cases because of the two different modes of calculation used, but the relative differences between the two techniques are similar showing the robustness of our analysis.

**Figure 6:**
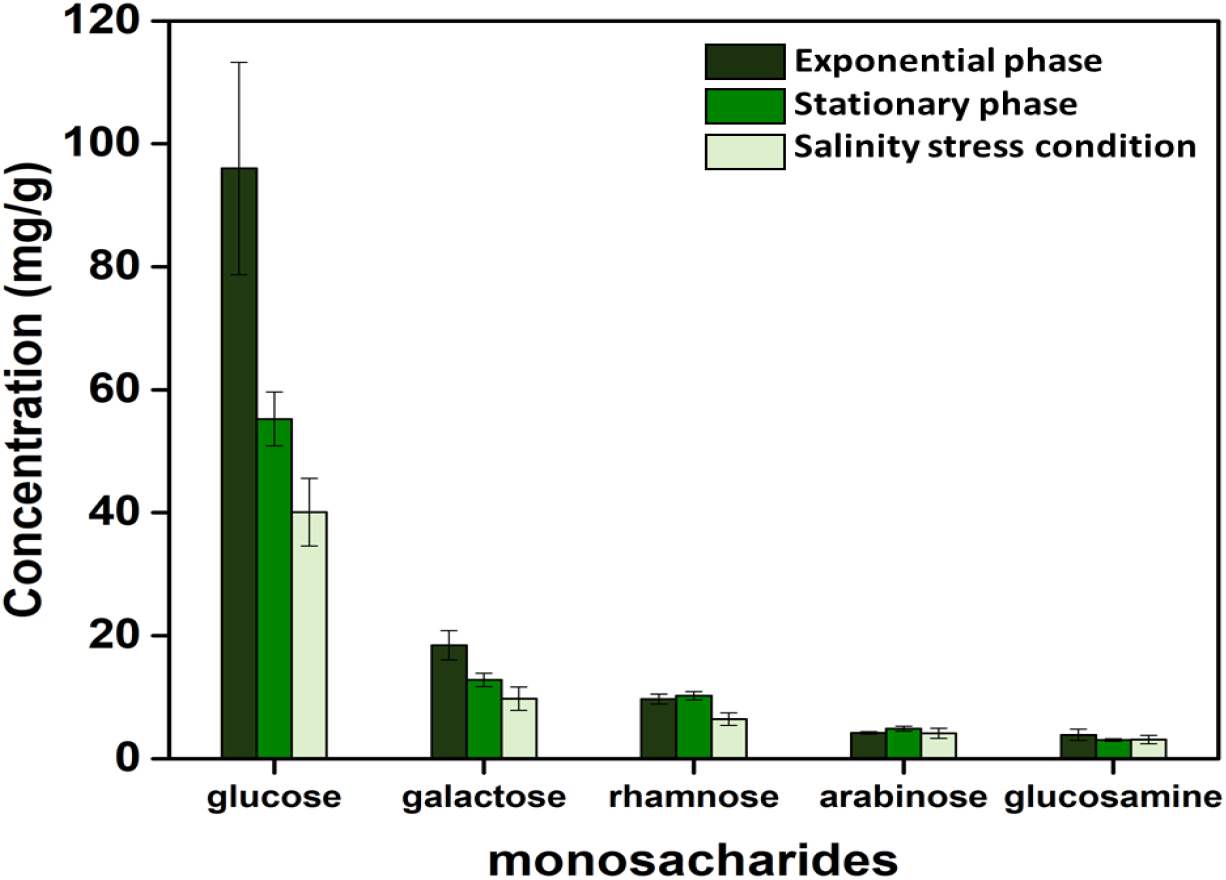
Monosaccharide composition of C. vulgaris cell wall in exponential phase, stationary phase and salinity stress condition (0.1M NaCl). The composition is expressed as milligram of monomer per gram of dry cell wall. The error bars indicate the deviation of the triplicates (n=3) from the average.

Different studies in the literature have analyzed the monosaccharidic composition of microalgae cell walls using similar experimental approaches. For example, in line with our findings, high glucose and galactose concentrations have also been reported by other authors in the case of *C. vulgaris* [9,50]. However, on the same microalgae species *C. vulgaris*, a study by Canelli *et al.* shows that the growth stage has no major effect on the monosaccharide composition [9]. This is different to what we observe here, however, in this study, a different strain of *C. vulgaris* was used, in different culture conditions. In the case of yeast microorganisms, it has been showed that the strains used, as well as their harvesting time play a critical role in the composition of the cell wall [51], thus the differences between our results and the study by Canelli *et al.* might be only due to these two parameters. This hypothesis is further comforted by a study performed on another microalgae species that shows that strain and culture conditions have an effect on the full biomass composition [52]. In addition, for different microalgae species, different compositions were reported at different growth phase. For example, major changes were observed in *Thalassiosira pseudonana*, where an increase in ribose, galactose, and mannose and a significant decrease of glucose [53] were reported for stationary phase cells. Additionally, in our case, cells in stationary phase and in saline stress conditions may have started consuming intercellular carbohydrate depositories, which could decrease the glucose fraction in the cell wall [9].

In these experiments, the polysaccharides present in the cell wall are hydrolyzed into monosaccharides. Identifying them however does not allow determining the polysaccharides they come from, which could be an interesting information to obtain if further cell wall degradations processes are developed to extract the cell’s contents. One way to identify the polysaccharides that could be present in the cell wall is to hydrolyze them with specific enzymes; if the substrate of the enzyme is present, the enzyme will release monosaccharides that can be detected by further HPLC analysis. So far, such studies have not been performed on microalgae cells walls, however, one study by Gerken *et al.* in 2013 consisted in treating cells on agar plates with specific enzymes; in this case an inhibition of cell growth was linked to the presence or not of their substrates in the cell wall [6]. Their work included the strain of *C. vulgaris* used in this study (strain CCAP211/11B); the main enzymes the authors identified as inhibiting cell growth for this strain are listed in Table 2. The polysaccharides that these enzymes hydrolyze and their corresponding monomers are also listed in this table. Hydrolysis with chitinase, chitosanase and lysozyme results in the release of the same monomer called glucosamine, which was identified in our experiments. This means that either chitin or chitosan (acetylated form of chitin) can be present in the cell wall of our strain. However, in Gerken’s study, chitosanase, which is specific to chitosan, only partially inhibited cell growth which could mean that perhaps chitin is the main form present in the cell wall of our cells. This could also mean that another type of glucosamine-based polymer is present: for instance, Canelli *et al.* stated that in stationary phase, a microfibrillar chitosan-like layer composed of glucosamine is present in *C. vulgaris* cell wall [9]. Lyzozyme can hydrolyze various substrates among which chitin and chitosan, but as stated in Gerken’s study, chitinase and lysozyme have different activities towards *C. vulgaris* cell wall, and the activity of lysozyme is required to expose other polymers in stationary phase cells. This thus may be in line with the findings of Canelli *et al.* and the glucosamine present in our cells, in stationary phase at least, could rather indicate the presence of another chitosan-like polymer. According to Gerken’s study, another enzyme that completely inhibited *C. vulgaris* cell growth is *β*-glucuronidase, however, we did not detect its corresponding monomer (glucuronic acid) in the cell wall. Either its concentration is lower than the limit of detection of our HPLC system, or the polysaccharides it hydrolyzes, glucuronides, are not present in our cells perhaps because of different culture conditions between Gerken’s and our study. Further, laminarinase, which can hydrolyze *β*-glucans into glucose monomers, has also been showed to completely inhibit *C. vulgaris* cell growth. Glucose is the most abundant monomer that we find in our experiments in all conditions. Thus this could mean that *β*-glucans form a large part of the polysaccharidic fraction of the cell wall, and the slight decrease in polysaccharides that we observe in stationary phase and in saline stress conditions compared to exponential phase (XPS data) may be due to the decrease in these conditions of these types of polysaccharides. Glucose is also the monomer constituting cellulose, however, Gerken *et al.* found that cellulase enzyme had no activity on *C. vulgaris* cells, meaning that this strain most likely does not contain cellulose, in contrast with other microalgae species [9,54]. Finally, the activity of pectinase and sulphatase enzymes found by Gerken *et al.*, could explain the presence of rhamnose, arabinose and galactose in our analysis, which could be part of pectic substances (rhamnose and arabinose) and of proteoglycans or glycosaminoglycans (galactose). Concerning the mannose and xylose that we found in our HPLC analysis, they could be coming from the degradation of mannoproteins or of hemicellulose. Despite the fact that some authors note the presence of hemicelluloses in cell wall of *Chlorella sp.*, there is no consensus on the type of hemicelluloses present in these cells [55].

Altogether, determining the saccharidic composition of the cell wall in this study allows to enhance our understanding of the cell wall. By comparing our data to the existing literature, we can make strong hypothesis on the polysaccharides present in the cell wall, which is an important point to understand the cell wall and develop strategies to disrupt it, for example through enzymatic degradations. Indeed, these disruption methods have been shown to consume less energy than mechanical or thermal treatments, and have already been successfully used for different types of microalgae [56]. In addition, as polysaccharides are often present directly at the cell surface, these hypothesis can also be important to understand the interacting behavior of cells, which can be determinant for example in harvesting processes using flocculation [21].

## CONCLUSIONS

In this work, three different techniques, AFM, XPS and chemical hydrolysis followed by HPAEC-PAD, were used to analyze *C. vulgaris* cell wall in different conditions relevant for the production of lipids used for biofuel production. The combination of these methods is original and has provided different information, which, taken together, has allowed to get new insights into the complexity of *C. vulgaris* cell wall and its dynamics depending on growth phase and culture conditions. For instance, we could show that in exponential phase, the cell wall is composed in similar proportions (approximately 40%) of polysaccharides (mainly glucose and galactose-based polysaccharides) and proteins and also contained around 20% of lipids. These proportions change with the growth phase; the cell wall evolves during growth and its composition changes with a large increase of lipids at the expanse of proteins. While the polysaccharidic fraction stays constant, the composition of this fraction also changes, with a decrease of glucose and galactose-based polymers. This composition variation is accompanied by an architectural changes that could be determined by probing the nanomechanical properties of the cell wall, which becomes significantly more rigid in stationary phase compared to exponential phase. Finally, when cells are submitted to a saline stress, their cell wall has a similar composition than for stationary phase cells but interestingly, it seems that the architecture of the cell wall is affected by the stress as they become a lot softer. Although in this case, the loss of turgor pressure induced by the osmotic stress may also partly explain the decrease in the rigidity observed. These new fundamental data, provided thanks to the original experimental approach developed in this study combining AFM, XPS and chemical hydrolysis, can be of great use to optimize important steps in microalgae-based biofuel production processes, such as harvesting or cell disruption. We believe these information will concretely contribute to the advancement of this field of research.

**Table 1:** Hypothesis on the polysaccharides present in C. vulgaris cell wall, formulated based on the enzymes that have been shown to inhibit cell growth by Gerken et al. [6]. The last column refers to references where it was found that the degradation of the corresponding polysaccharides led to the release of the monomers indicated.

| Enzyme [6] | Polysaccharide | Monomer | Reference |
| --- | --- | --- | --- |
| **Chitinase**<br>(+++) | chitin | N-acetyl-D-glucosamine<br>(GlcNAc) | [57] |
| **Chitosanase**<br>(++) | chitosan<br>oligosaccharides | β-(1,4)-D glucosamine | [58] |
| **Lysozyme**<br>(+++) | Peptidoglycan<br>lipoproteins,<br>lipopolysaccharides<br>(LPS), and some<br>hydrophobic peptides | N-acetylglucosamine N-acetyl muramic acid | [59,60] |
| **B-Glucuronidase**<br>(+++) | Glucuronide/ β-<br>glucuronides | Glucuronic Acid | [61] |
| **Laminarinase**<br>(+++) | β-glucans | Glucose | [62] |
| **Pectinase**<br>(+++) | Pectic substances;<br>pectin, protopectin and<br>pectic acids | Rhamnose<br>arabinan, galactan or<br>arabinogalactan | [63] |
| **Sulphatase**<br>(+++) | Proteoglycans,<br>glycosaminoglycans | galactose | [64,65] |

## ACKNOWLEDGMENTS

C. F.-D. is a researcher at CNRS. C. F.-D. acknowledges financial support for this work from the Agence Nationale de la Recherche, JCJC project FLOTALG (ANR-18-CE43-0001-01). The authors would like to thank Pauline Herviou for her help with HPLC measurements, and Malak Souad Ftouhi for help with microalgae cultures.

## CONFLICT OF INTEREST

The authors declare no conflicts of interest.

## Notes

### Competing Interest Statement

The authors have declared no competing interest.

