## Supplementary for "Combining AFM, XPS and chemical hydrolysis to understand the complexity and dynamics of *C. vulgaris* cell wall composition and architecture"

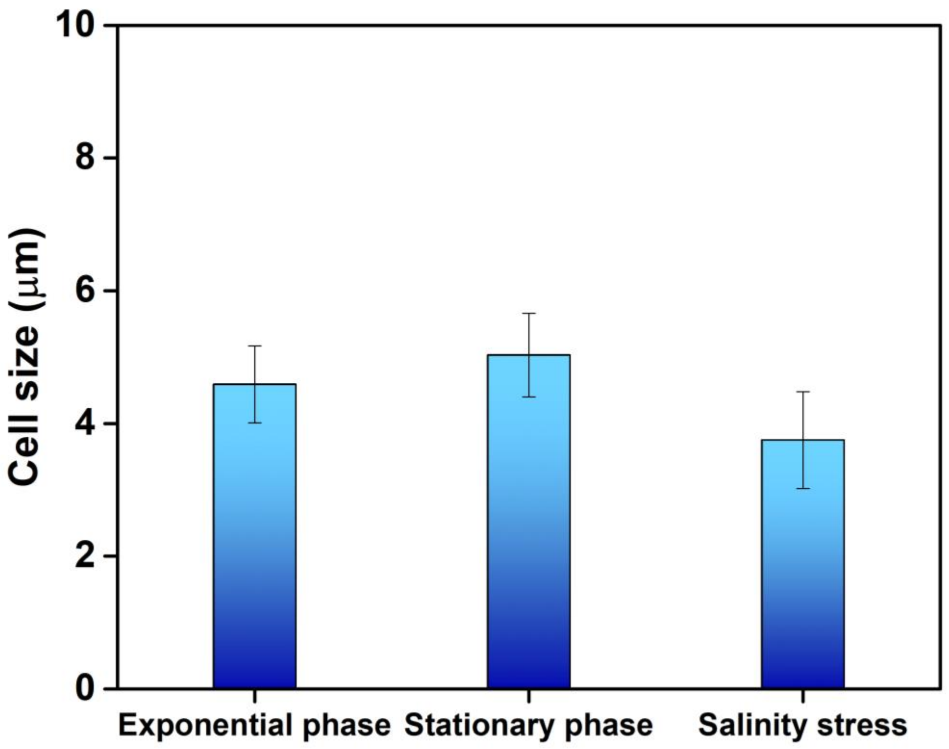


**Supplementary Figure S1.** *C. vulgaris* cell size in exponential phase, stationary phase and salinity stress condition (0.1M NaCl). The error bars indicate the deviation of the triplicates (n=3).

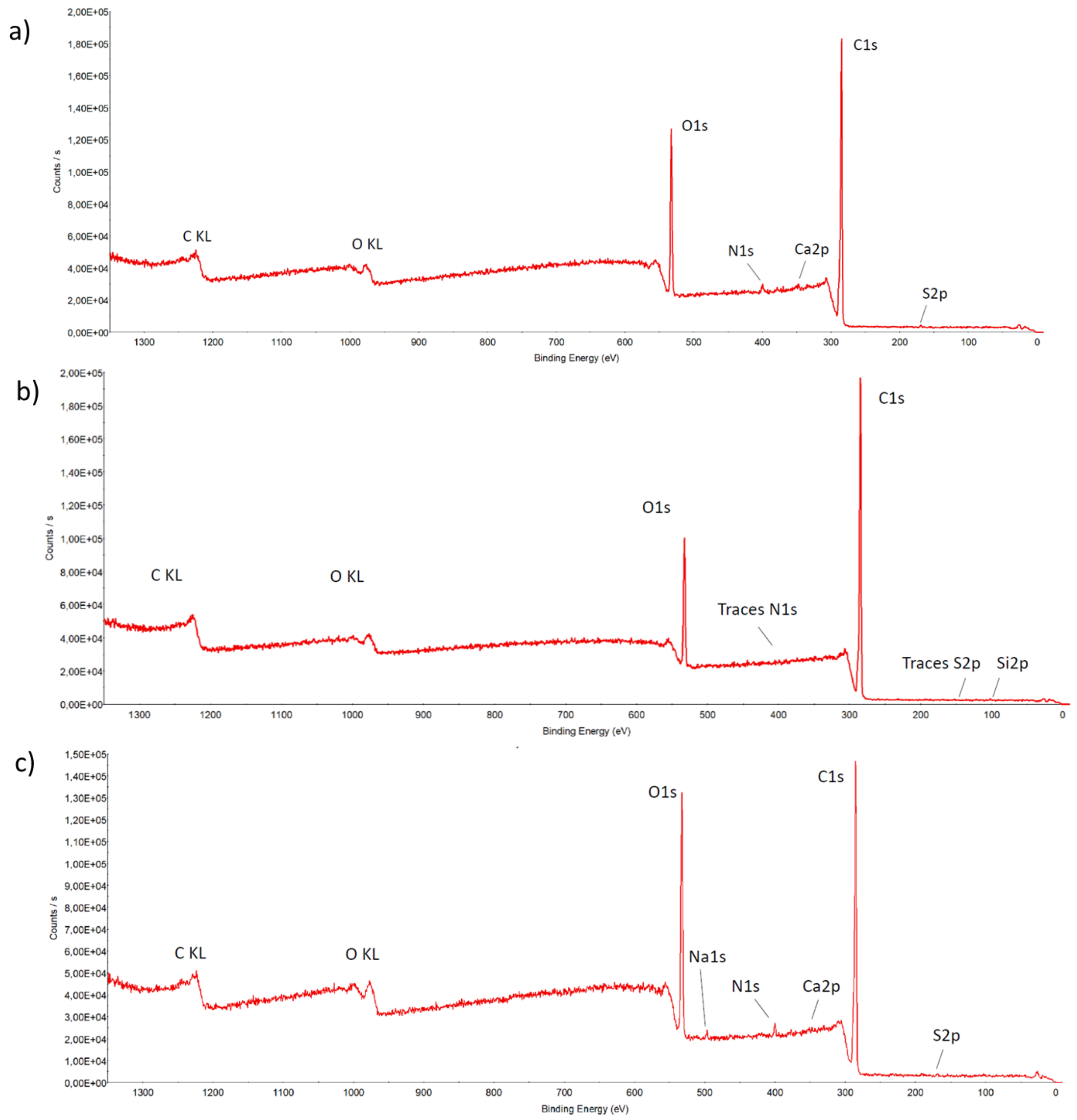


**Supplementary Figure S2.** XPS overall spectra recorded on *C. vulgaris* cell walls isolated from cells grown in a) exponential phase, b) stationary phase, and c) salinity stress condition.

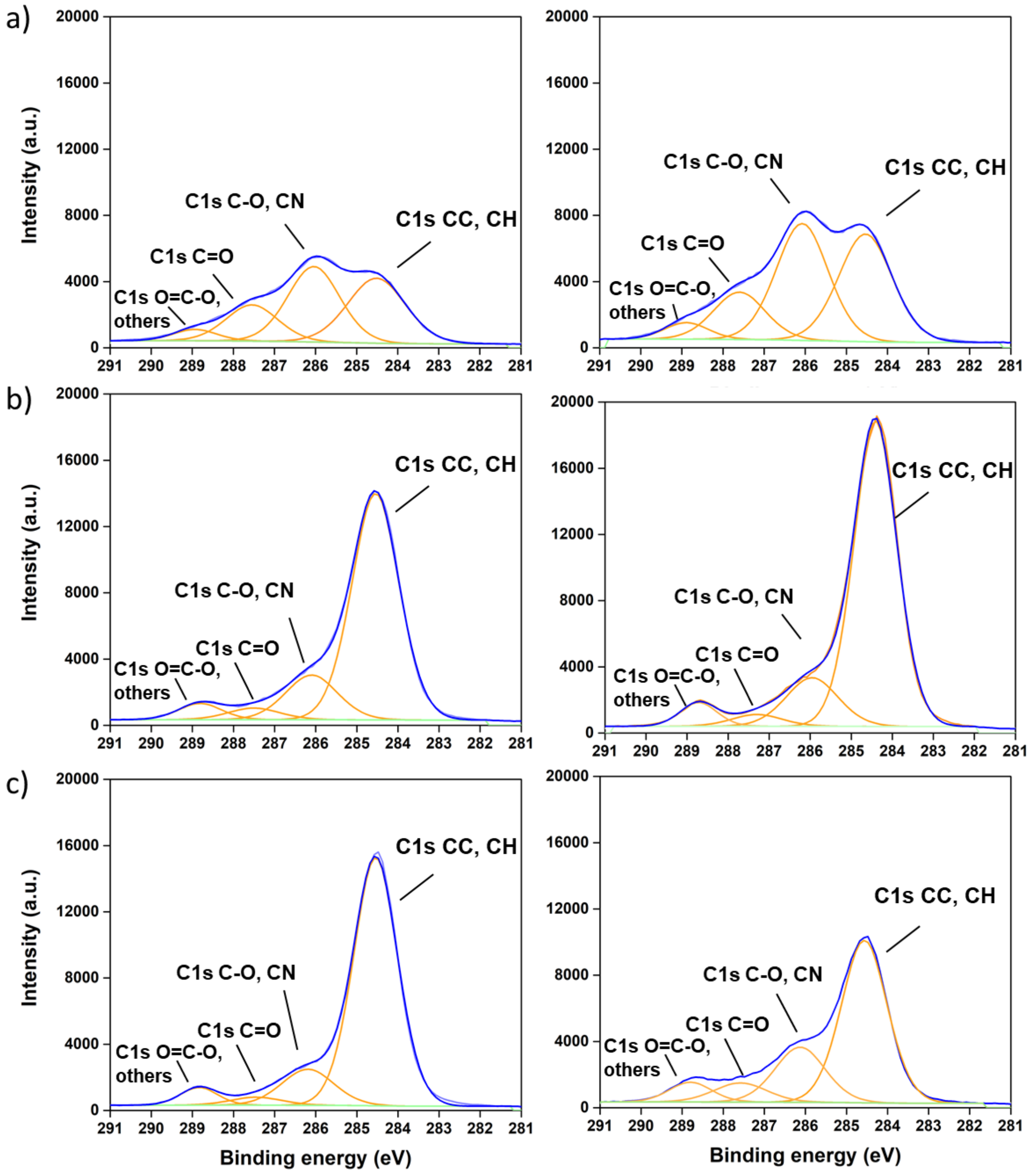


**Supplementary Figure S3.** XPS Carbon 1s peaks recorded on *C. vulgaris* cell walls isolated from cells grown in a) exponential phase, b) stationary phase and c) salinity stress condition. Average atomic percentages and standard deviations are presented in Supplementary table 1-3 from triplicates (n=3).

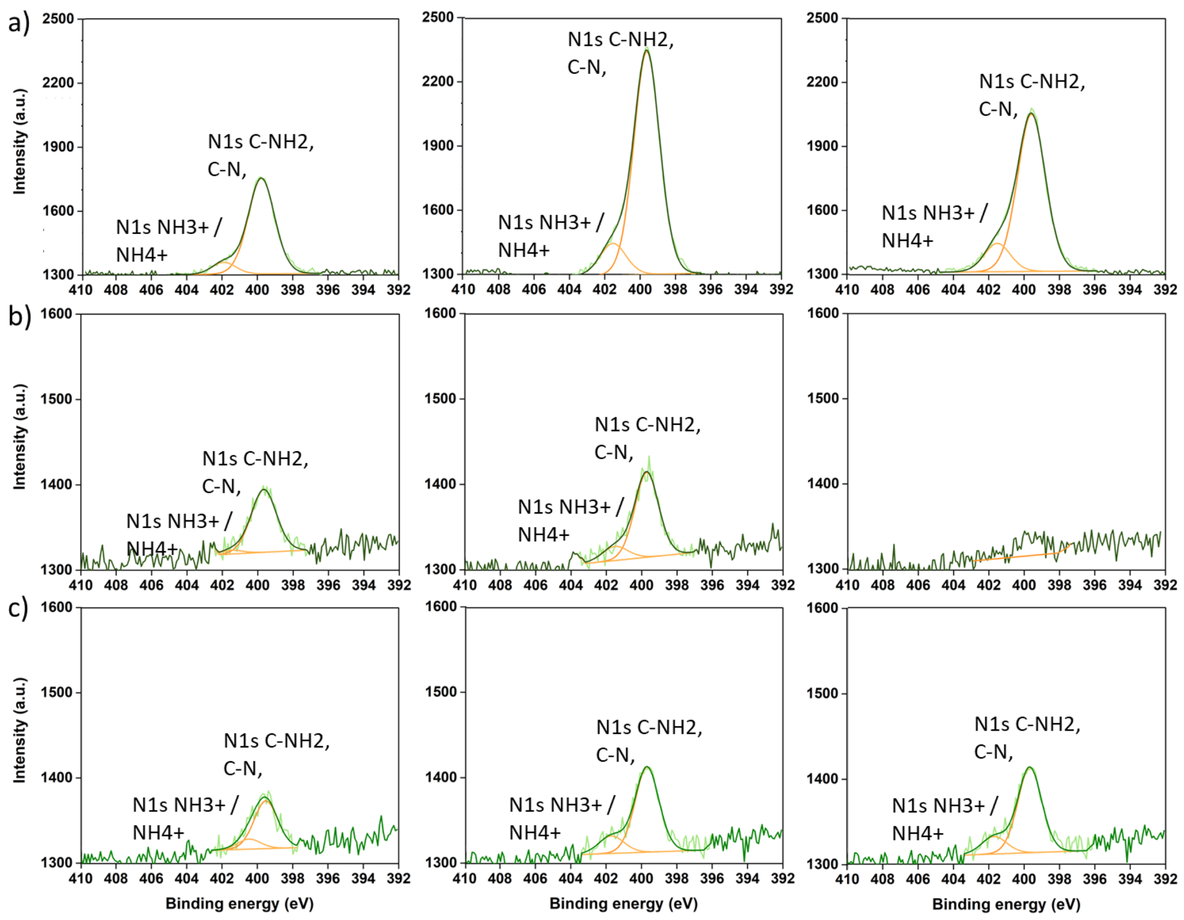


**Supplementary Figure S4.** XPS Nitrogen 1s peaks recorded on *C. vulgaris* cell walls isolated from cells grown in a) exponential phase, b) stationary phase and c) salinity stress condition. Average atomic percentages and standard deviations are presented in Supplementary table 1-3 from triplicates (n=3).

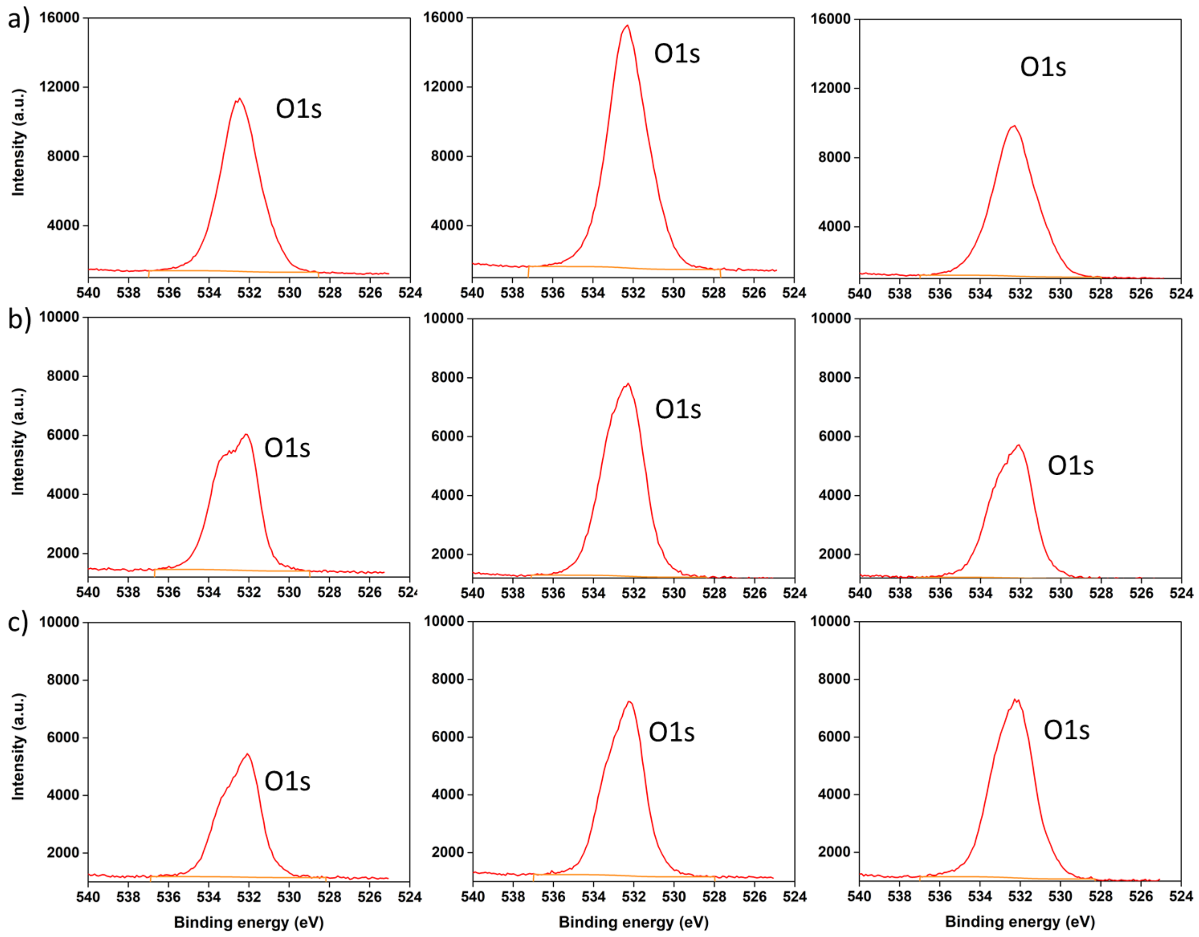


**Supplementary Figure S5.** Oxygen 1s peaks recorded on *C. vulgaris* cell walls isolated from cells grown in a) exponential phase, b) stationary phase and c) salinity stress condition. Average atomic percentages and standard deviations are presented in Supplementary table 1-3 from triplicates (n=3).

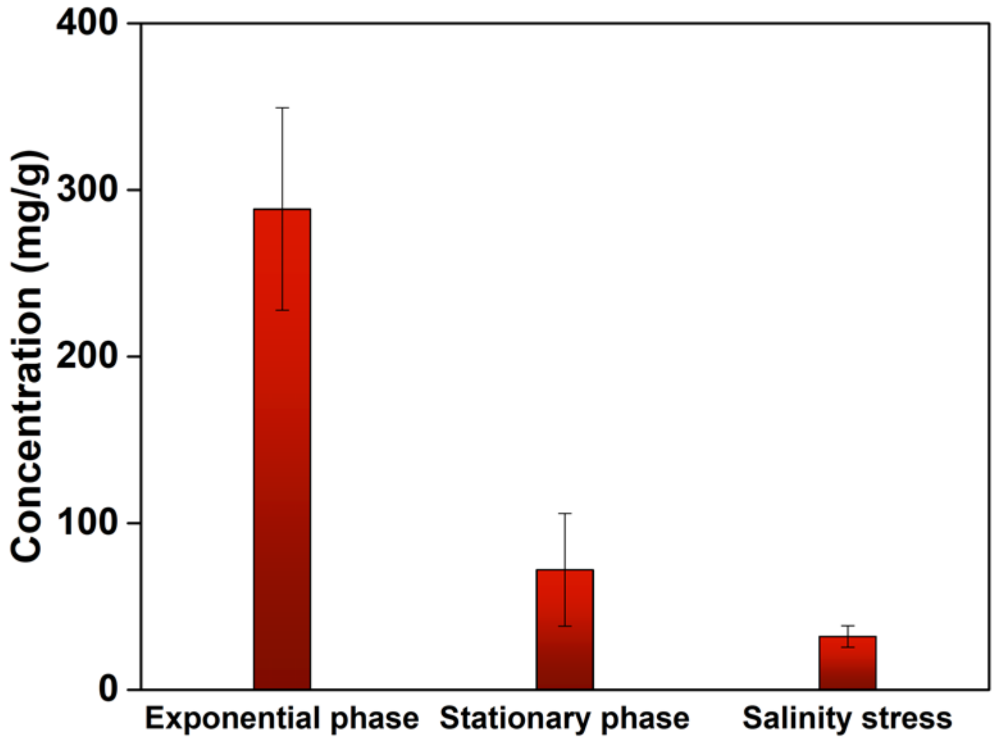


**Supplementary Figure S6.** Protein concentration of *C. vulgaris* cell wall in exponential phase, stationary phase and salinity stress condition (0.1M NaCl). The composition is expressed as miligram of protein per gram dry cell wall. The error bars indicate the deviation of the triplicates (n=3) from the average.

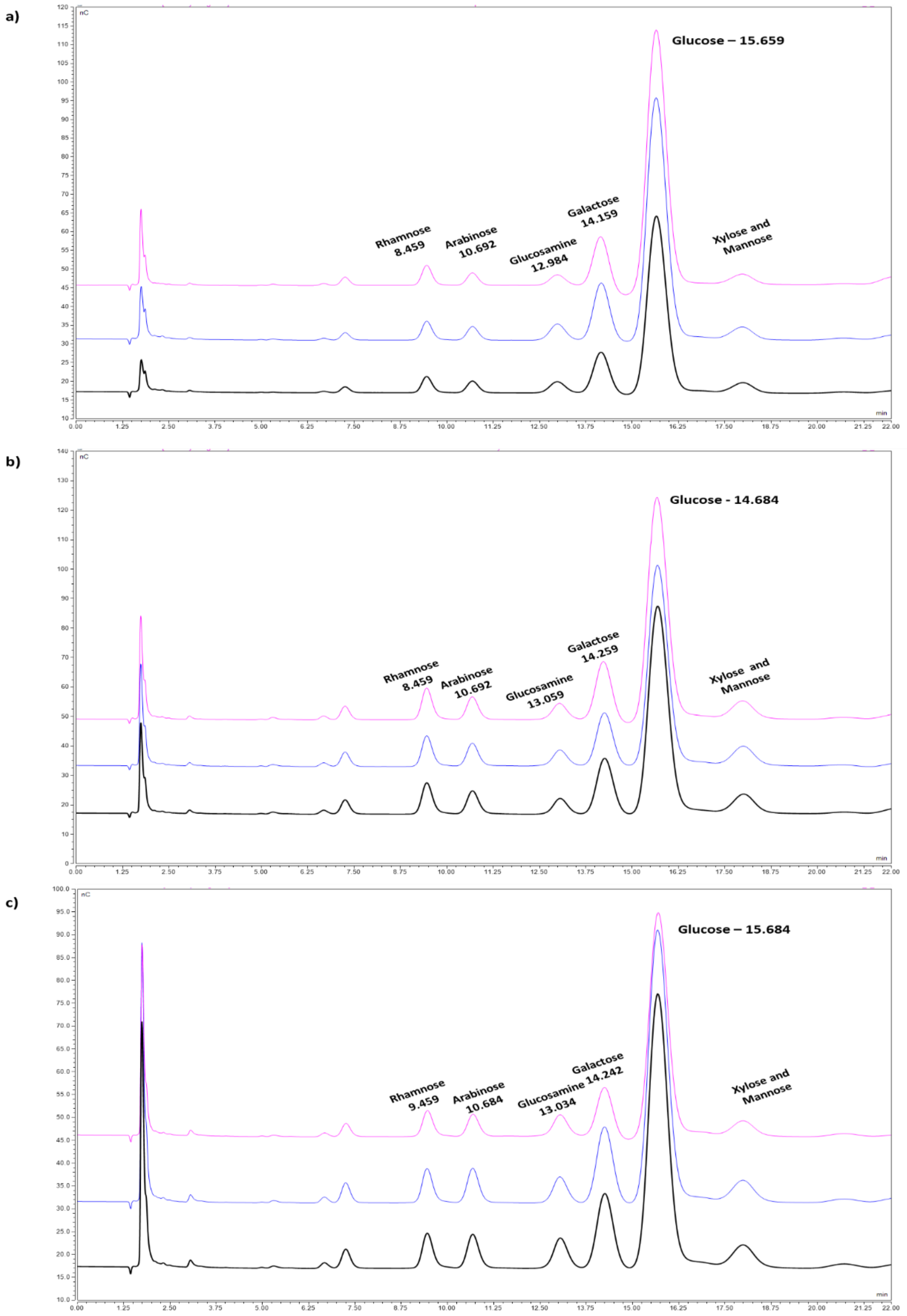


**Supplementary Figure S7.** HPAEC-PAD chromatograms of the monosaccharide composition of the *C. vulgaris* cell wall in a) exponential phase b) stationary phase and c) salinity stress condition. Different colors show the biological replicates.

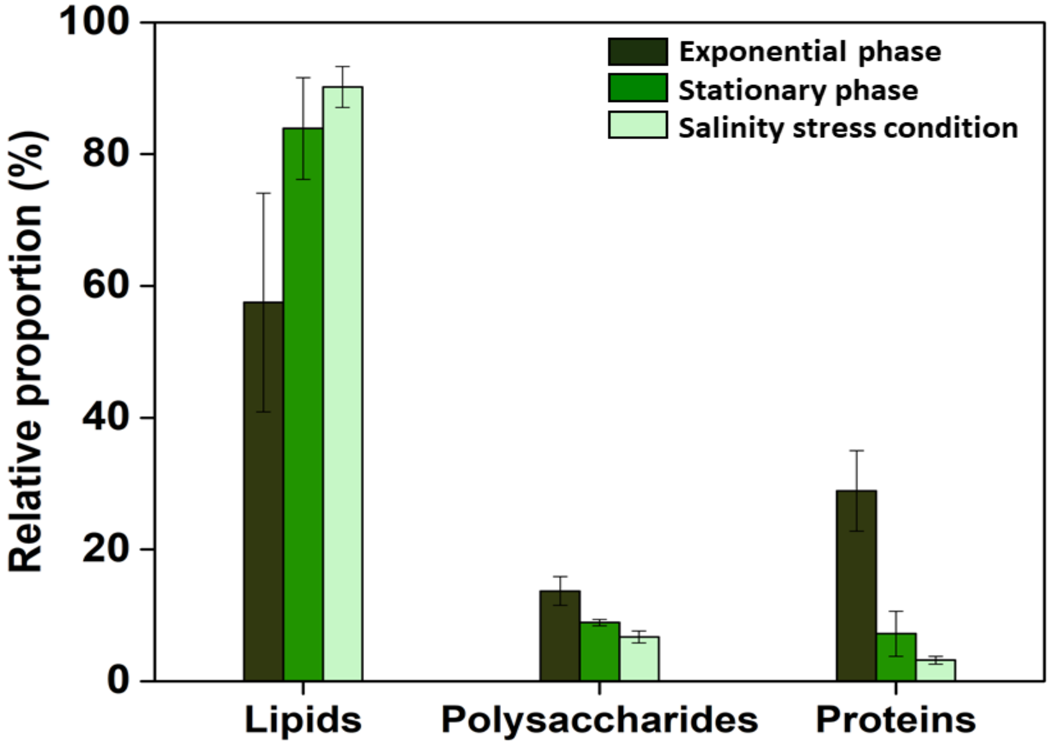


**Supplementary Figure S8.** Biochemical composition of *C. vulgaris* cell wall in exponential phase, stationary phase and salinity stress condition (0.1M NaCl). Polysaccharide compositions were based on HPLC analysis whereas protein compositions were based on bicinchoninic acid (BCA) protein assay kit and lipid composition were obtained subtracting the protein and polysaccharides amount from the total cell wall. The composition is expressed as percentage. The error bars indicate the deviation of the triplicates (n=3).

**Supplementary Table S1.** XPS atomic percentages obtained for the fitted components for cells in exponential phase. The values correspond to the average and standard deviations obtained from triplicates.

| Name | Peak BE (eV) | Atomic % |
|---|---|---|
| **Si2p** | 102.1 ± 0.0 | 0.5 ± 0.1 |
| **S2p S2p3/2 -SH (thiol)** | 163.5 ± 0.1 | 0.0 ± 0.0 |
| **S2p S2p1/2 -SH (thiol)** | 164.7 ± 0.1 | 0.0 ± 0.0 |
| **S2p S2p3/2 O-SO3- (sulfate)** | 168.3 ± 0.1 | 0.4 ± 0.0 |
| **S2p S2p1/2 O-SO2- (sulfate)** | 169.5 ± 0.1 | 0.2 ± 0.0 |
| **C1s CC, CH** | 284.5 ± 0.0 | 28.2 ± 6.4 |
| **C1s C-O, CN** | 286.1 ± 0.1 | 23.4 ± 2.2 |
| **C1s C=O, O-C-O** | 287.6 ± 0.0 | 10.0 ± 1.6 |
| **C1s O=C-O, others** | 288.9 ± 0.1 | 3.6 ± 0.8 |
| **K2p** | 292.7 ± 0.1 | 0.3 ± 0.0 |
| **Ca2p** | 347.1 ± 0.1 | 0.6 ± 0.1 |
| **N1s C-NH2, C-N** | 399.7 ± 0.1 | 2.5 ± 0.7 |
| **N1s NH3+ / NH4+** | 401.7 ± 0.2 | 0.4 ± 0.1 |
| **O1s** | 532.4 ± 0.1 | 30.1 ± 2.5 |
| **Na1s** | / | / |

**Supplementary Table S2**. XPS atomic percentages obtained for the fitted components for cells in stationary phase. The values correspond to the average and standard deviations obtained from triplicates.

| Name | Peak BE (eV) | Atomic % |
|---|---|---|
| **Si2p** | 102.1 ± 0.0 | 0.2 ± 0.0 |
| **S2p S2p3/2 -SH (thiol)** | / | / |
| **S2p S2p1/2 -SH (thiol)** | / | / |
| **S2p S2p3/2 O-SO3- (sulfate)** | 168.3 ± 0.0 | 0.1 ± 0.0 |
| **S2p S2p1/2 O-SO2- (sulfate)** | 169.5 ± 0.0 | 0.0 ± 0.0 |
| **C1s CC, CH** | 284.6 ± 0.0 | 60.2 ± 6.4 |
| **C1s C-O, CN** | 286.1 ± 0.0 | 14.9 ± 2.4 |
| **C1s C=O, O-C-O** | 287.5 ± 0.0 | 3.8 ± 0.8 |
| **C1s O=C-O, others** | 288.8 ± 0.0 | 4.5 ± 0.4 |
| **K2p** | / | / |
| **Ca2p** | 347.0 ± 0.0 | 0.2 ± 0.0 |
| **N1s C-NH2, C-N** | 399.7 ± 0.0 | 0.3 ± 0.0 |
| **N1s NH3+ / NH4+** | 401.5 ± 0.0 | 0.0 ± 0.0 |
| **O1s** | 532.4 ± 0.1 | 16.1 ± 2.8 |
| **Na1s** | / | / |

**Supplementary Table S3.** XPS atomic percentages obtained for the fitted components for cells in saline stress conditions. The values correspond to the average and standard deviations obtained from triplicates.

| Name | Peak BE (eV) | Atomic % |
|---|---|---|
| **Si2p** | 101.9 ± 0.0 | 0.7 ± 0.0 |
| **S2p S2p3/2 -SH (thiol)** | 163.0 ± 0.0 | 0.1 ± 0.0 |
| **S2p S2p1/2 -SH (thiol)** | 164.2 ± 0.0 | 0.1 ± 0.0 |
| **S2p S2p3/2 O-SO3- (sulfate)** | 168.1 ± 0.1 | 0.1 ± 0.0 |
| **S2p S2p1/2 O-SO2- (sulfate)** | 169.3 ± 0.1 | 0.1 ± 0.0 |
| **C1s CC, CH** | 284.6 ± 0.0 | 56.8 ± 8.2 |
| **C1s C-O, CN** | 286.2 ± 0.1 | 14.6 ± 2.5 |
| **C1s C=O, O-C-O** | 287.5 ± 0.1 | 4.4 ± 1.6 |
| **C1s O=C-O, others** | 288.8 ± 0.0 | 4.7 ± 0.5 |
| **K2p** | 292.2 ± 0.0 | 0.1 ± 0.0 |
| **Ca2p** | 346.7 ± 0.0 | 0.2 ± 0.0 |
| **N1s C-NH2, C-N** | 399.5 ± 0.1 | 0.5 ± 0.3 |
| **N1s NH3+ / NH4+** | 401.3 ± 0.6 | 0.1 ± 0.0 |
| **O1s** | 532.3 ± 0.0 | 18.2 ± 3.5 |
| **Na1s** | 1070.0 ± 0.0 | 0.2 ± 0.0 |

**Supplementary Table S4.** Relative biomass proportions of lipids, proteins and polysaccharides in C. vulgaris cell wall in the different conditions used in this study. The errors indicate the deviation from the average of the triplicates (n=3) recorded in two different zones in samples coming from two independent cultures.

| | **Exponential phase** | **Stationary phase** | **Salinity stress** |
|---|---|---|---|
| **Lipids** | 18.9 ± 9.3% | 57.2 ± 11.1% | 60.5 ± 7.0% |
| **Proteins** | 41.4 ± 9.2% | 11.8 ± 8.7% | 6.2 ± 2.2% |
| **Polysaccharides** | 39.7 ± 3.5% | 31.0 ± 2.5% | 33.2 ± 5.0% |
